# Dyslexic children show altered temporal structure of the nonlinear VEP

**DOI:** 10.1101/697201

**Authors:** Sheila Crewther, Jacqueline Rutkowski, David Crewther

**Affiliations:** School of Psychological Science, La Trobe University, Victoria 3086, Australia; Brain Sciences Institute, Swinburne University of Technology, Victoria 3122, Australia

## Abstract

The neural basis of dyslexia remains unresolved, despite many theories relating dyslexia to dysfunction in visual magnocellular and auditory temporal processing, cerebellar dysfunction, attentional deficits, as well as excessive neural noise. Recent research identifies perceptual speed as a common factor, integrating several of these systems. Optimal perceptual speed invokes transient attention as a necessary component, and change detection in gap paradigm tasks is impaired in those with dyslexia. This research has also identified an overall better change detection for targets presented in the upper compared with lower visual fields. Despite the magnocellular visual pathway being implicated in the aetiology of dyslexia over 30 years ago, objective physiological measures have been lacking. Thus, we employed nonlinear visual evoked potential (VEP) techniques which generate second order kernel terms specific for magno and parvocellular processing as a means to assessing the physiological status of poor readers (PR, n=12) compared with good readers (GR, n=16) selected from children with a mean age of 10yr. The first and second order Wiener kernels using multifocal VEP were recorded from a 4° foveal stimulus patch as well as for upper and lower visual field peripheral arcs. Foveal responses showed little difference between GR and PR for low contrast stimulation, except for the second slice of the second order kernel where lower peak amplitudes were recorded for PR *vs* GR. At high contrast, there was a trend to smaller first order kernel amplitudes for short latency peaks of the PR *vs* GR. In addition, there were significant latency differences for the first negativity in the first two slices of the second order kernel. In terms of peripheral stimulation, lower visual field response amplitudes were larger compared with upper visual field responses, for both PR and GR. A trend to larger second/first order ratio for magnocellularly driven responses suggests the possibility of lesser neural efficiency in the periphery for the PR compared with the GR. Stronger lower field peripheral response may relate to better upper visual field change detection performance when target visibility is controlled through flicking masks. In conclusion, early cortical magnocellular processing at low contrast was normal in those with dyslexia, while cortical activity related to parvocellular afferents was reduced. In addition, the study demonstrated a physiological basis for upper versus lower visual field differences related to magnocellular function.

## Introduction

Developmental Dyslexia (DD) is an ‘impairment in the acquisition of literacy skills despite normal intelligence, socioeconomic and educational circumstances’ in (DSM-IV) (American Psychiatric Association, 2000), and now placed with specific learning disorder in DSM-5 (American Psychiatric Association, 2013). It affects approximately 5-10% of school-aged children (Habib, 2000) with slow reading and poor spelling typically retained in adults who suffered from DD (Snowling, 2000;Winner et al., 2001). Activation of the left angular gyrus forms a meta-analytical focus in fMRI studies of those with DD (Temple, 2002;Temple et al., 2003) - conforming with the belief that the primary deficit in dyslexia involves phonological processing.

The rapid activation of visual mechanisms supporting object recognition coupled with oculomotor control and superior integration of visual and auditory with phonological processing are all necessary for fluent reading and continue to attract research into the fundamental neural mechanisms of dyslexia. As such, the magnocellular visual system is thought to provide many of the aspects of parietal visual processing as well as shifts of visual attention (reviewed in (Stein and Walsh, 1997;Habib, 2000)) involved in reading. Indeed, Stein and Walsh (1997) proposed an abnormality in the magnocellular visual pathway as a cause of dyslexia. This is empirically supported by experiments showing that DD children possess a deficit in the processing of low spatial frequency and luminance contrast stimuli and lowered temporal visibility from as early as 1980 (Lovegrove et al., 1980). Lowered motion and motion coherence sensitivity (Cornelissen et al., 1995;Talcott et al., 2000), as well as reduced brain activation in Area V5/hMT+ to moving stimuli (Eden et al., 1996;Demb et al., 1997;1998) are also characteristic of those with dyslexia. However, the relation between reading performance and motion processing has been somewhat contentious.

Parietal cortex transient attentional processes have been implicated in the aetiology of dyslexia (Steinman et al., 1998;Facoetti et al., 2000;Bosse et al., 2007). Rutkowski et al. (2003) using a gap paradigm change detection task, identified the sudden appearance of the second stimulus, coupled with circular place holders, as a potential driving factor. This study showed that a much longer initial presentation time was needed in a group of dyslexic children in order to just detect change. This finding fits with a parietal locus, shown in fMRI experiments (Beck et al., 2001;Beck et al., 2006) and causally through transcranial magnetic stimulation (TMS) (Tseng et al., 2010). Also, magnetoencephalography (MEG) has been used to show that the ability to shift attention is the likely function at play (Pammer et al., 2006;Pammer, 2013). The literature on change detection also implicates visual masking as making a contribution to change blindness (Rensink, 2002). Pertinent to this finding, Crewther et al. (2019) showed that metacontrast masking (with dynamic random dot annuli around each item of the change detection stimuli) provides a measure of change detection performance with higher contrast masking causing greater distraction, as evidence by poorer performance. These metacontrast change detection studies showed superior change detection for upper visual field locations.

Skottun (2000) rightly points out that the magnocellular pathway and the dorsal cortical stream are different entities (Skottun, 2015), but fails to acknowledge the dominance of magnocellular inputs into early dorsal cortical stream. Logically, this demands the use of physiological techniques to detect where in the brains of those with reading problems, lie the differences in activity from those who read normally.

Thus we aimed to measure physiological aspects of the magno (M) and parvocellular (P) processing of good and poor readers through analysis of the nonlinearities of the multifocal visual evoked potential (Klistorner et al., 1997;Crewther et al., 1999). This technique has been used in the analysis of M and P associations with autistic tendency (Sutherland and Crewther, 2010;Jackson et al., 2013), with the relation between magnocellular efficiency and flicker fusion in normals (Brown et al., 2018b) and in magnocellular performance in ageing (Brown et al., 2018a). The ability to record evoked responses from stimuli in different parts of visual field simultaneously, allows us to measure M and P processing of stimuli in upper and lower visual field regions, and hence investigate the neural mechanism of an upper visual field advantage reported in the change detection of letter stimuli (Rutkowski et al., 2003).

## Methods

### Participants and screening

29 participants (age range 7-12 yr) were recruited from a larger investigation across schools and summer learning camps. The Institutional Ethics Committee approved the study and informed consent was obtained from parents before testing commenced with any of the children. Children were screened for visual abnormalities and were excluded if any uncorrected binocular or refractive errors were present. The Neale Analysis of Reading Ability (NARA)(Neale, 1997) was used to assess dyslexia based on published norms (Neale, 1999). The NARA has elements of word recognition, comprehension and reading speed, and has been the basis for more recent reading tests (Spooner et al., 2004). Non-verbal intelligence was assessed using the Raven’s Coloured Progressive Matrices Test (RCPM) (Raven et al., 1990). Assignment of participants to poor reading (PR) (chronological-reading age ≥1.5 yr) or good reading (GR) groups was on the basis of two criteria: chronological age compared with reading age, with normal level of intelligence. One participant was excluded on the basis of Raven’s score (see Table 1).

**Table 1:** Mean Chronological Age and Mean Reading Age (NARA) ± 1 SE for the Poor and Good readers.

| Reading | N Rows | Chronological Age<br>(yr) | Reading Age<br>(yr) |
| --- | --- | --- | --- |
| PR | 12 | 10.21±0.44 | 7.75±0.39 |
| GR | 16 | 10.38±0.35 | 10.78±0.43 |

### mfVEP

One of the advantages of multifocal visual evoked potentials (mfVEP) is that evoked responses from separate parts of the visual field can be collected simultaneously – the full decorrelation of stimulus sequences ensuring independence. In this way we could study both foveal processing and peripheral processing at the one time (Sutter, 2000).

### Stimuli

A close-packed array of 37 hexagons (central patch plus 3 rings of 6, 12 and 18 hexagonal patches respectively) was employed. The foveal stimulus subtended 4° for participants at 45cm distance with a total subtense of 30° of visual field. Background luminance was 50 cd/m^2^. In addition, the summed responses of the upper and lower halves, excluding the foveal patch (with eccentricity from 2° −20°) were analysed in order to extend earlier reports of upper versus lower visual field change detection performance differences (Rutkowski et al., 2003). Recordings were made for two temporal contrast levels – 24% and 96%. These values were chosen as those that gave the greatest separation between second order kernel contrast responses functions (see Klistorner et al. (1997))

**Figure 1:**
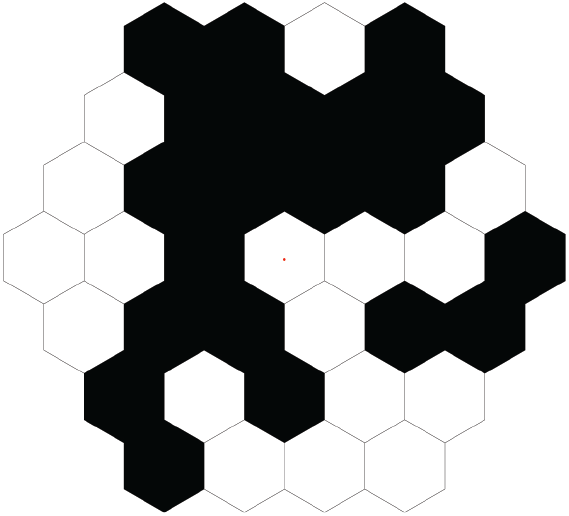
Hexagonal array with 37 patches used for the mfVEP recording. Each patch was stimulated with a pseudo random decorrelated sequence

The stimulus sequence employed was a m=14 binary pseudorandom m-sequence presented using a VERIS system (EDI, USA), with each patch operating a pseudo-random sequence decorrelated from all others. The frame rate of the Apple Mac computer monitor was 67 Hz. The four-minute recording period was broken into 4 epochs to ensure a relaxed state in the participants. The first order kernels and the first two slices of the second order (assessing the effect of prior stimuli on the VEP at varying time scales) were analysed. Peak latencies and amplitudes were extracted using time windows designed to capture individual minima or maxima around the population mean for all participants.

## Results

### mfVEP

The VEP first order kernels and the first two slices of the second order in response to the central, foveal stimulus, are shown in separate figures for the poor reader (PR) and good reader (GR) groups at low contrast (Fig.2) and at high contrast (Fig.3).

**Figure 2:**
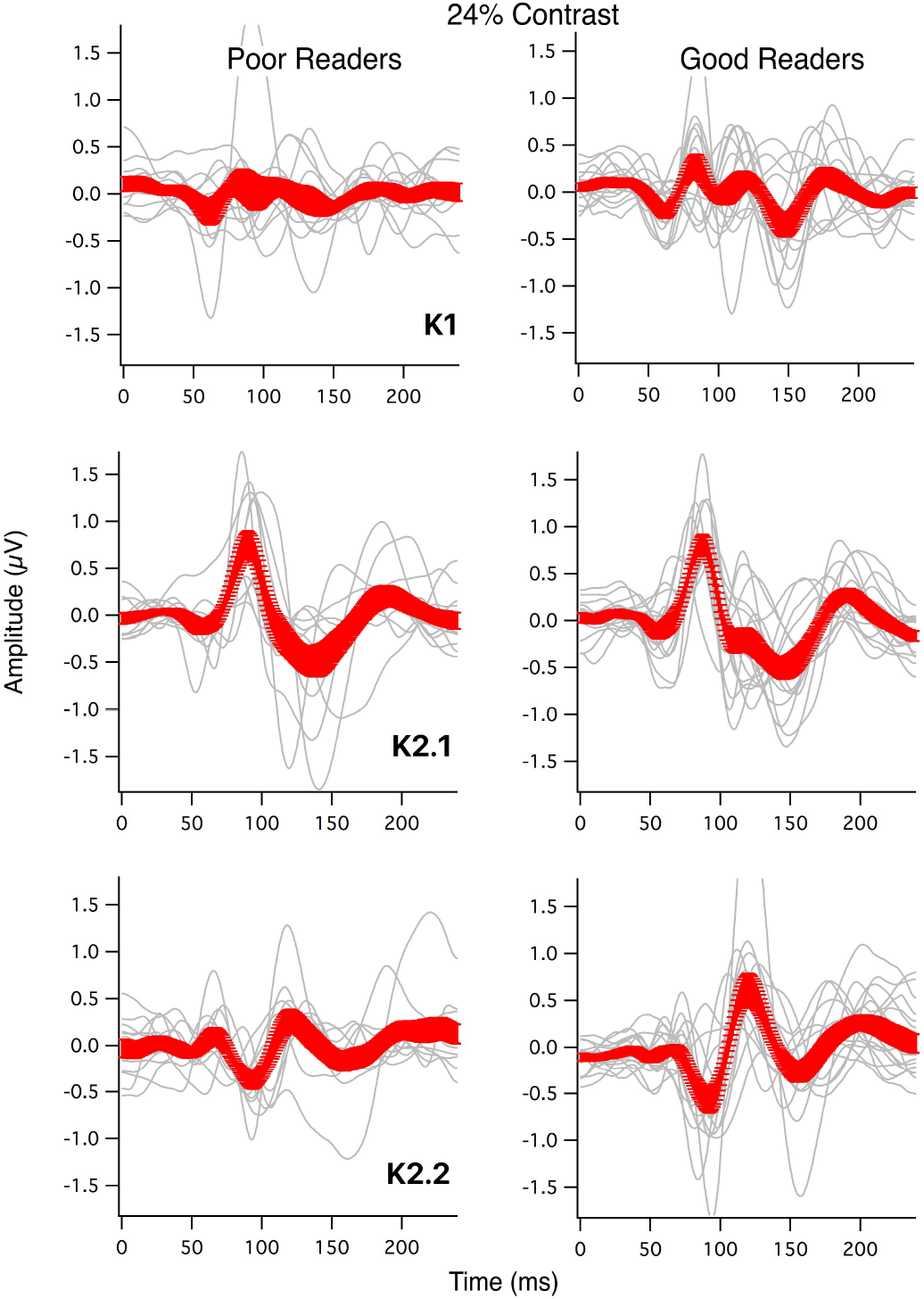
Individual response kernels for poor readers (PR) and Good Readers (GR) under low contrast (24% temporal contrast) stimulation. The mean waves ±1SE are shown in red.

**Figure 3:**
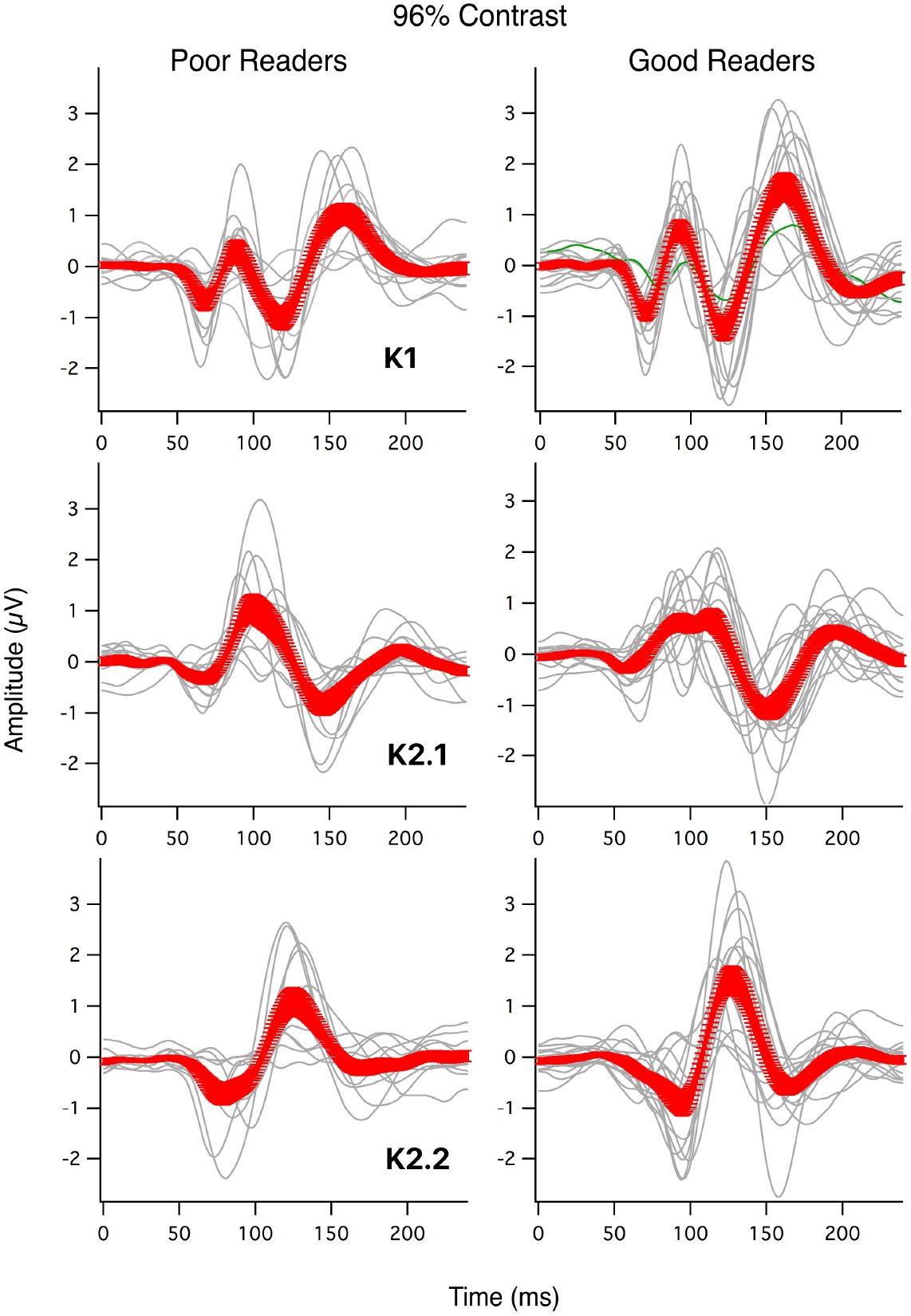
Individual response kernels for poor readers (PR) and Good Readers (GR) under high contrast (96% temporal contrast) stimulation. The mean waves ±1SE are shown in red.

Overall, with low contrast (24%) stimulation, the mean waveform for the PR group showed small deflections in the first order response, comparable to the level of noise. However, the first slice of the second order responses (largely magnocellularly generated (Klistorner et al., 1997;Jackson et al., 2013)) were well defined for both PR and GR groups and very similar in magnitude and waveform. The second slice of the second order responses was reduced in magnitude for major N95P120 peak with smaller amplitude for the PR group (t-ratio = 2.025, df=25.55, Prob >t = 0.0272).

At high contrast (96% modulation), the amplitudes of the peaks were typically larger for the NR compared with the PR groups, through just outside significance (see Figs. 3 and 4). The latency of the N1 peak (around 75ms for the K2.2 kernel slice) was significantly different between the two reading groups (*p*<.02). While there was no evidence of magnocellular abnormality in the low contrast VEP K2.1 kernels, it is apparent from Figure 3 that for high contrast stimulation there is an abnormal positivity around P100. Closer inspection of the individual waves contributed by the GR and PR demonstrate the existence of a secondary positivity or perhaps an extended width K2.1 P100 peak in the NR compared with the PR as observed in young adults with high autistic tendency (Sutherland and Crewther, 2010).

**Figure 4:**
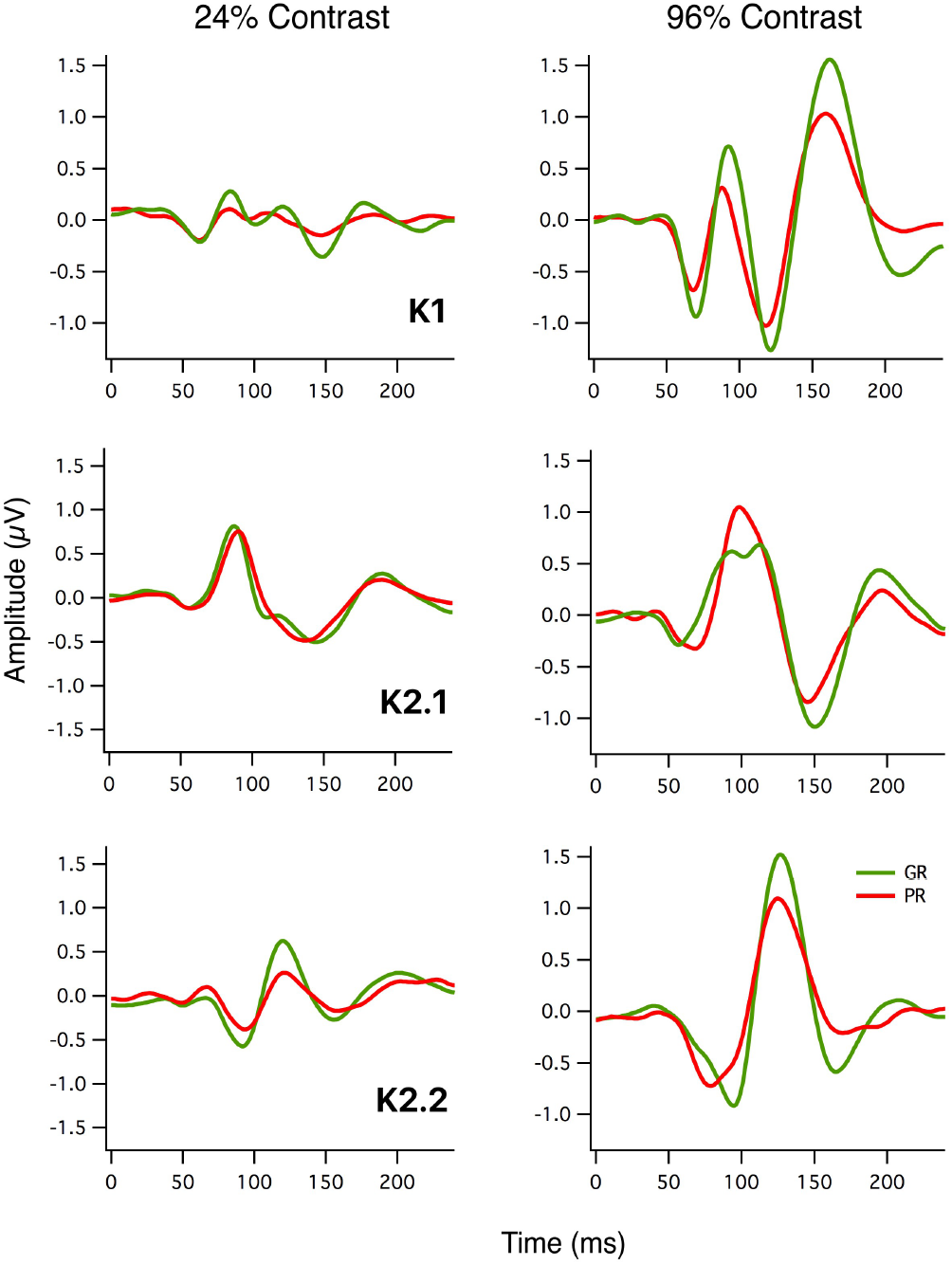
First and second order (K2.1, K2.2) mfVEP kernel responses recorded at 24% and 96% contrast (foveal stimulus) for good (GR – green lines) and poor readers (PR – red lines).

Comparing the first and second order kernel waveforms of the PR and GR (Figure 4), several observations are clear. The nonlinear peak amplitude associated with M pathway processing are not different between PR and GR, and normal in shape for low contrast stimulation. Secondly, there are systematic reduction in wave amplitudes for PR compared with GR, for those peaks associated with P pathway processing (Klistorner et al., 1997;Jackson et al., 2013).

On the basis of earlier psychophysical results indicating an upper visual field advantage for change detection (Rutkowski et al., 2002;2003) evidence was sought for UVF versus LVF physiological response differences. Inspection of the individual responses at peripheral location it was clear that less signal to noise was obtained as one stimulated into the periphery. Hence the responses for the 15 upper visual field (U) and 15 lower visual field (L) hexagonal patches (see Figure 1) were summed and compared for the GR and PR. As is clear from Figure 5, the major peak (P90) of the first order kernel responses recorded from LVF showed greater mean amplitudes compared with those for UVF in both the GR and PR groups, with the statistical comparison marginally significant (Paired t-test, df = 27, *p*=.05).

**Figure 5:**
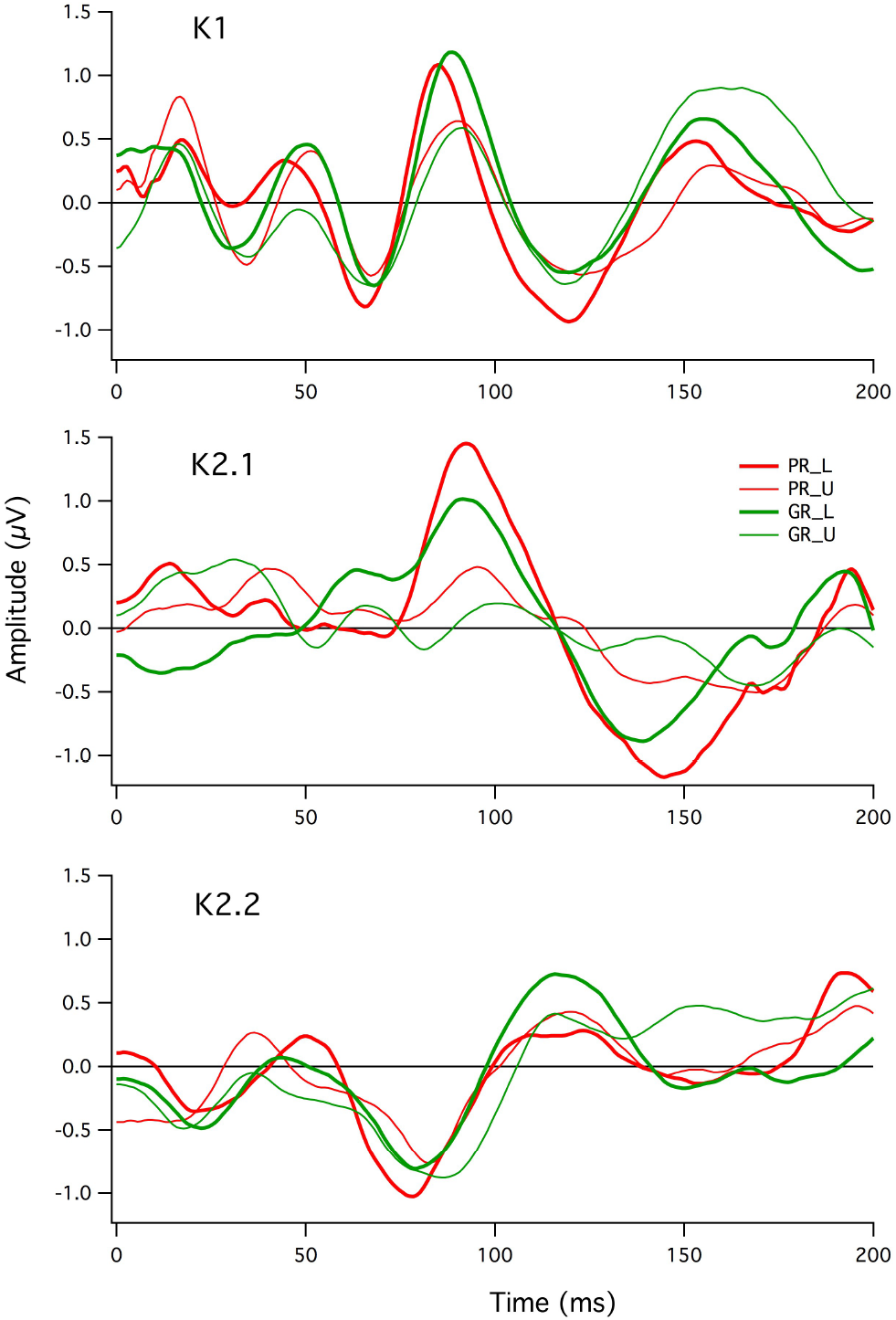
Comparison of mean Upper (U) (thin traces) and Lower (L) (bold traces) visual field responses for first order kernel (K1) and the first and second slices of the second order kernel (K2.1, K2.2) for good (GR green lines) and poor (PR red lines) readers. There is a clear lower visual field enhancement in terms of peak amplitudes in the first order (K1) and first slice of the second order (K2.1) waveforms.

The first slice of the second order response (K2.1) shows a stronger visual field effect than does the first order response(repeated measures ANOVA *F*(1,26) = 9.682, *p*<.004), however, although there is a qualitatively greater amplitude in K2.1 for the PR group compared with NR group, there was no significant interaction between Visual Field and Reading Group (GR, PR).

## Discussion

The mfVEP recordings presented here give a quite clear indication that magnocellular non-linearities (K2.1 major peak – see Fig 2) are not dissimilar between GR and PR at low contrast. However, waveform shape differences were observed at high contrast, similar in nature to the K2.1 P100 disturbance for young adults with high autistic tendency, while using similar nonlinear VEP techniques (Sutherland and Crewther, 2010;Jackson et al., 2013). Interestingly, it was the PR group that appeared to show a simpler waveform, possibly reflecting a lesser complexity of processing connectivity *cf* GR (Paulesu et al., 1996). Until the VEP developmental trajectory of M and P mechanisms (Crewther et al., 1999) are more fully understood, the interpretation of this difference will remain somewhat opaque.

Differences were manifest between PR and GR groups for low contrast stimulation in terms of the K2.2 N90P125 peak. This parvocellularly generated nonlinearity (Jackson et al., 2013) is smaller for the PR compared with the GR. This P-generated reduction is present at trend level across all of the kernel slices evaluated, both in first order N110P160 amplitude of the first order (K1) and in the N90P130 amplitude of the K2.2 second slice. In addition, there was a significant latency differences of the earliest negative deflection of the K2.2 slice. This finding does not exclude the involvement of magnocellular processing in the etiology of dyslexia – it just appears that when taken in isolation from the other kernel elements, the M-generated K2.1 nonlinearities are similar in amplitude. On closer inspection it appears that a pattern of smaller K1 N60P90 peaks coupled with equal or larger amplitude K2.1 peaks is emerging. Thus, the results are consistent with poorer neural efficiency of cortical responses of the M pathway, though larger sample sizes are required to confirm this notion.

The necessity of transient attention, generally thought to apply through the intraparietal cortex area of the dorsal stream, for change detection, and the significantly poorer change detection performance by PR compared with GR indicates that the other sources of transient information reaching parietal cortex need to be studied. The significance of the difference in change detection reported between good and poor readers (Rutkowski et al., 2003) suggests that rapidity of recognition might play an important role in the development of fluent reading (see Grosser and Trzeciak (1981), and this is borne out by experiments associating perceptual speed with reading ability (McLean et al., 2011). The notion that cortical oscillations need to be synchronised in order for the reading process to be efficient (Pammer, 2013;Goswami et al., 2014) has brought confusion about the direction of causality into mechanisms of dyslexia, with some opinion pointing to sensory differences in dyslexia arising because of the difficulty in reading – i.e. a reversal of the normal causality (see the different points of view:(Goswami, 2015b;a;Lobier and Valdois, 2015)). The current paper regarding early cortical physiology, and another parallel one on rapid attention measured though change detection in a gap paradigm (Crewther et al., 2019) add significantly to this debate.

In terms of upper versus lower visual field stimulation, the clear bias for lower visual field VEP responses at first seems at odds with the reported poorer LVF vs UVF change detection (Rutkowski et al., 2003). However, reflection on the nature of the distraction provided by the dynamic mask flicker as introduced by Crewther et al. (2019) or the sudden appearance of a high contrast static frame surrounding letters in the second stimulus in the gap paradigm used (Rutkowski et al., 2003) suggests that the effect of masking may be greater in the lower visual field precisely because of the increase in magnocellular sensitivity there.

## Acknowledgements

The authors would like to acknowledge support from the philanthropic Andrew Fildes Foundation for sponsoring the summer camps, and the Australian Research Council for funding a discovery project.

